# A common deliberative process underlies model-based planning and patient intertemporal choice

**DOI:** 10.1101/499707

**Authors:** Lindsay E. Hunter, Aaron M. Bornstein, Catherine A. Hartley

**Author notes:** These authors contributed equally to this work (Lindsay E. Hunter, Aaron M. Bornstein).

## Abstract

Humans and animals consistently forego, or “discount” future rewards in favor of more proximal, but less valuable, options. This behavior is often thought of in terms of a failure of “self-control”, a lack of inhibition when considering the possibility of immediate gratification. However, rather than overweighting the near-term reward, the same behavior can result from failing to properly consider the far-off reward. The capacity to plan for future gains is a core construct in Reinforcement Learning (RL), known as “model-based” planning. Both discounting and model-based planning have been shown to track everyday behaviors from diet to exercise habits to drug abuse. Here, we show that these two capacities are related via a common mechanism – people who are more likely to deliberate about future reward in an intertemporal choice task, as indicated by the time they spend considering the choice, are also more likely to make multi-step plans for reward in a sequential reinforcement learning task. In contrast, the degree to which people’s intertemporal choices were driven by a more automatic bias did not correspond to their planning tendency, and neither did the more standard measure of discounting behavior. These results suggest that the standard behavioral economic measure of discounting is more fruitfully understood by decomposing it into constituent parts, and that only one of these parts corresponds to the sort of multi-step thinking needed to make plans for the future.

## 1 Introduction

The virtue of patience is more than a cliché. In the lab, patience is often measured as the degree to which participants are willing to forego a reward of given value in exchange for eventual delivery of a reward of greater value. According to the standard framework, the value of the delayed reward is “discounted” or diminished in proportion to the time one must wait to receive it. The rate at which future rewards are discounted as a function of time (“discount rate”) provides an idiosyncratic metric of relative patience (“shallow” discounting) vs. impulsivity (“steep” discounting). Critically, this measure has been shown to have considerable trait-like stability across behavioral domains that may reflect relative prioritization of long-term reinforcement, including academic performance, drug abuse, diet and exercise habits [1, 2, 3].

A longstanding puzzle is exactly how this richly predictive and otherwise stable trait can reliably be altered sometimes dramatically by attributes of the decision setting or internal state of the decision-maker [4]. Answering this question will require understanding the cognitive and neural mechanisms involved in carrying out intertemporal choices. Although the framework of temporal discounting provides a straightforward, economical metric for assessing individual differences in delay discounting, it does not speak to the underlying cognitive mechanisms through which subjective intertemporal preferences arise [5]. Traditionally, both economists and psychologists have framed discounting as a phenomenon of self-control [6], with shallow discounting reflecting the inability to resist the hedonic pull of immediate reward. In these terms, “impulsive” behavior arises from hypersensitivity to immediate rewards, decreased cognitive control, or a combination of both [7]. However, a preference for sooner reward can arise either from over-valuation of relatively small quantities on the immediate horizon, or under-valuation of relatively larger rewards to be received later. Recent work has called attention to this latter phenomenon, highlighting the role of the prospective valuation process in patient intertemporal choice [8, 9]. For instance, Peters & Bchel (2010) found that cueing personal future events reduced discounting of rewards to be delivered at the time of those events, and that neural evidence of episodic imagery in response to these cues predicted the extent of this effect.

Formalizing the idea that intertemporal choice characteristics arise from prospective valuation, Gabaix & Laibson (2017) [10] recently showed that the appearance of time preferences can emerge in an agent who has no inherent preference over time per se, but who instead simply relies on forecasts of the subjective utility of consuming a future reward (by imagining, e.g., a future vacation to Philadelphia). Such an agent can exhibit discounting as long as they are “myopic” that is, as long as the reliability of their forecasts decreases with the amount of time remaining until the future consumption event. One candidate mechanism for myopic forecasting is sequential evidence accumulation, such as captured by the drift-diffusion model (DDM; [11]). Suggestively, performance in tasks that are well-described by this mechanism is susceptible to the same sorts of framing manipulations e.g. attention allocation, time pressure, and cognitive load [12, 13, 14] that are known to affect intertemporal choice, and the DDM and related models have been successfully applied to model the dynamics of intertemporal choice [15, 16, 17].

In the present study, we hypothesized that this sort of mechanism, when parameterized such that deliberation scales with the time until receipt of reward, might reproduce choice patterns in a standard intertemporal choice task and correspond to a separate, external measure of prospective valuation namely, “model-based” planning in sequential reinforcement learning. Model-based RL refers a set of decision strategies that use the contingency structure of the environment in order to make choices that maximize long-run gain. The computational distinction between model-based RL and its “model-free” counterpart, which makes choices on the basis of direct experience with stimulus-reward associations, has proven a fruitful framework for understanding many features of value-based choice [18]. Empirically, a link between discounting and model-based planning is implied by the fact that prospective valuation is thought to underlie both forms of behavior [19, 20]. This symmetry is complemented by experimental evidence linking model-based learning to behaviors associated with self-control, such as compulsion disorders [21], and by the fact that manipulations that affect discounting and evidence accumulation also affect model-based planning [22]. The prospect of identifying an explicit link between these measures is appealing because it offers formal computational basis for understanding the mechanisms through which prospective deliberation influences action evaluation. Just as reliable forecasts of future reward facilitate patience in intertemporal choice, model-based reinforcement learning promotes long-term reward maximization by simulating potential outcomes [18].

**Figure 1:**
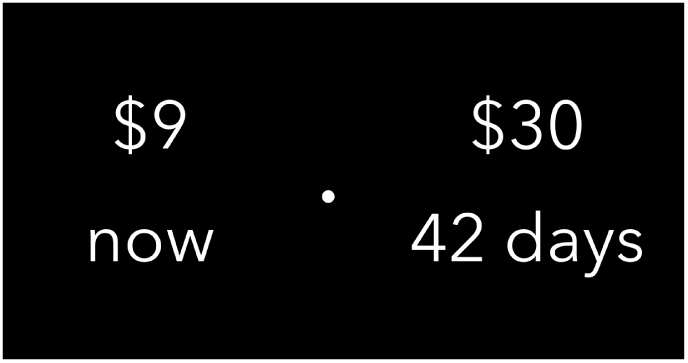
Intertemporal choice task. Participants made a series of 102 binary choices between a reward ($1-$85) to be delivered immediately following the experiment (smaller-sooner, SS), and a larger reward ($10-$95) to be delivered between four and 180 days in the future (larger-later, LL) via time-locked debit card. Participants had six seconds to make a choice; their response time was recorded along with their decision.

Here, participants performed two experimental tasks. In the first task, a large group of participants made several independent choices between smaller, immediate rewards and larger, later rewards(Figure 1. A subset of these participants also performed a sequential “two-step” choice task designed to index model-based choice behavior [23, 24]. We first used a hyperbolic model to estimate individual discount rates, a measure of their relative patience, from participants choices in the intertemporal choice task. We further leveraged information about decision time to decompose choice into deliberative and reflexive components. Specifically, we fit choices and reaction times with a sequential sampling model that separately measured both an individuals overall bias towards choosing the patient option as well as the degree to which they deliberated about each choice. This model both reproduced the choice patterns fit by the hyperbolic model and provided a mechanistic explanation for the relationship between patience, choice variability, and decision time. Importantly, both the bias and deliberative components of the model independently tracked an individuals degree of patient choice. We next derived, from the sequential RL task, an index reflecting the extent to which each participants behavior reflected model-based planning. We found that the deliberative component of the sequential sampling model tracked individuals tendency to use model-based planning, while their discount rate and bias did not. This finding suggests that both kinds of choices rely on a common prospective evaluation mechanism, the decomposition of which can be used to better understand the patterns and predictiveness of preferences in intertemporal choice.

## 2 Methods

563 participants (ages 18–66) recruited at New York University gave written informed consent following New York University Committee on Activities Involving Human Subjects (UCAIHS) and were compensated and debriefed at the end of the session. On the basis of behavior in the Intertemporal Choice Task, 31 participants were excluded for a bias of greater than 95% towards choosing the LL or SS options. 7 additional participants were excluded for completing fewer than 96 trials of the ITC task. A further 63 participants were excluded for a fitted hyperbolic discount rate that was outside the range reliably estimable by our choice set (see Intertemporal Choice Task below for further details). The remaining 462 (291 female; mean age = 22.86 years, SD = 5.95; age and sex for nine participants are omitted from these statistics due to data loss) participants were included in the reported analysis.

**Figure 2:**
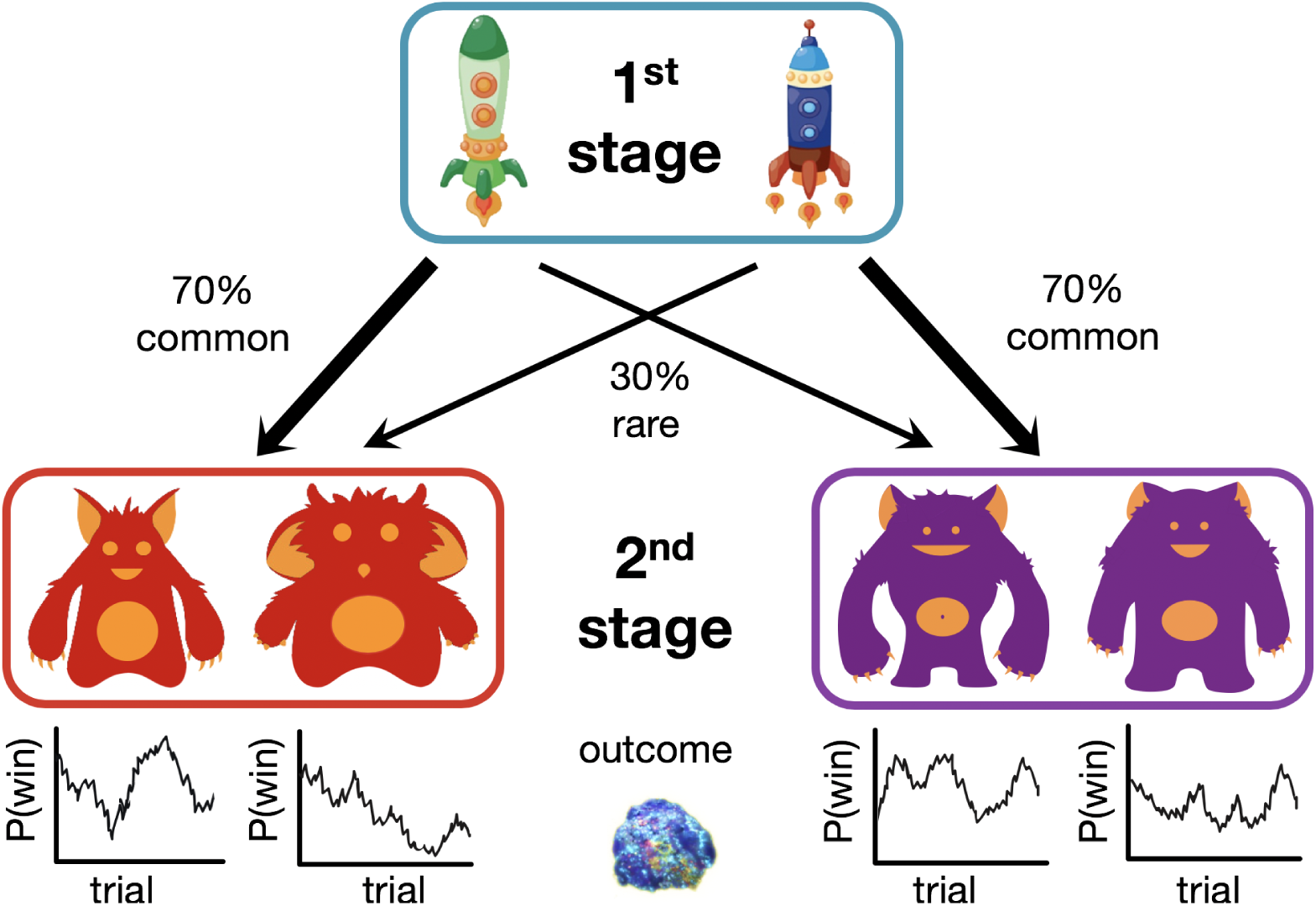
Two-step Reinforcement Learning Task. For each trial, subjects made a first-stage choice between two spaceships, each of which led more frequently (70% versus 30%) to one planet than the other. For example, choosing the blue spaceship traveled to the red planet with 70% probability (common transition), and traveled to the purple planet with 30% probability (rare transition). At this second stage, subjects chose between two aliens and were rewarded (a picture of space treasure) or not (an empty circle) according to a slowly drifting probability which was independent for each alien.

44 participants (ages 18–26) recruited at Weill Cornell Medical College completed the ITC task as well as a second decision-making task, the Two-step Reinforcement Learning Task (Daw et al., 2011). These participants gave written informed consent and debriefed at the end of the session. To control for any ordering effects (e.g., fatigue) the sequence of tasks was counterbalanced across participants. 5 participants were excluded for a bias of greater than 95% towards choosing the LL or SS options. A further 7 participants were excluded on the basis of a discount rate outside the range estimable by our choice set. The remaining 32 volunteers (17 female; mean age = 22.7 years, SD = 2.50) were included in the reported analyses.

Because the ITC task was identical for each participant population, we combined these data for analysis, giving N=494 total participants.

### 2.1 Behavioral Tasks

#### 2.1.1 Intertemporal Choice Task

Participants used key presses to make a series of 102 binary choices between a smaller amount of money they would receive immediately (same-day) and a larger amount of money that would be paid after a variable number of days 1. Across trials, the magnitude of monetary amounts ranged from $1 to $85 for the smaller, sooner (SS) payment, and $10 to $95 for the larger, later (LL). The SS payment was delivered at the end of the session (0 days), while the LL was delivered between four and 180 days in the future. The side (left vs. right) on which the smaller-sooner (SS) vs. larger-later (LL) reward appeared was counterbalanced across trials. After the stimuli were presented, participants had 6s to make a selection by pressing either “1”- or “0”- keys which corresponded to the left-and right- rewards respectively. After the choice was entered, a checkmark appeared indicating the chosen option for 500ms. There was a variable inter-trial-interval (ITI) of 3–5s. Since prior work suggests that real and hypothetical choices may fundamentally differ [25], participants were paid for one randomly selected trial from the full set of decisions and instructed to make choices according to their true preferences. Bonuses were awarded in the form of an Amazon gift card sent electronically via email the day of the experiment. The balance and activation date of the gift card were matched to the magnitude and delay associated with the chosen option on the selected trial. This payment policy was designed to minimize potential choice biases arising due to any increased uncertainty associated with receipt of delayed rewards.

#### 2.1.2 Two-step Reinforcement Learning Task

To capture individual differences in goal-directed choice we used an adapted version of a two-step Reinforcement Learning paradigm designed to dissociate model-based and model-free learning strategies (as in (Decker et al., 2016); adapted from Daw et al., 2011). Before starting the task, all participants completed an interactive tutorial and verbally confirmed their understanding of the instructions.

The primary objective of the task was to collect as much space treasure as possible. At the beginning of each trial (Figure 2), participants made a first-stage choice between a blue spaceship and a green spaceship, which was followed by a probabilistic transition (rare 30%, common 70%) to one of two 2nd planets (red vs. purple). The blue spaceship commonly (70%) transitioned the red planet and rarely (30%) transitioned to the purple planet whereas the red spaceship commonly (70%) transitioned the blue planet and rarely (30%) transitioned to the red planet. Subjects were informed that the transitions from first stage choices (i.e., spaceships) to second stage states (i.e., planets) were symmetric and remained fixed throughout the task. At the 2nd stage state, subjects made a choice between one of two aliens and were rewarded (a picture of space treasure) or not (an empty circle) according to a slowly drifting probability which was independent for each alien. Reward probabilities changed slowly, stochastically, and independently (independent Gaussian random walks bound between .2 and .8) over the course of the task, incentivizing exploration of 2nd stage stimuli (aliens) throughout the experiment. All choices were subject to a 3s response deadline. Once a response was submitted, an animation highlighted the chosen stimulus (spaceship or alien) for 1s, as well as the outcome (space treasure or an empty circle) for 1s after 2nd stage choices. If participants failed to make a first- or 2nd stage choice, both stimuli were covered by a red X for 3s before the next trial started. Participants completed 200 trials separated by a 1s inter-trial interval.

The fixed probabilistic transition between 1st and 2nd stage states enabled the distinction of model-based and model-free choices by examining the influence of the previous trial on the subsequent first-stage choice. A model-free learner is likely to repeat a previously rewarded first-stage choice (“stay”), regardless of the transition type that led to the reward (a main effect of reward on subsequent first-stage choices). In contrast, a model-based chooser considers the transition structure, reflected by an interaction effect of transition type (common vs. rare) and reward on “stay or switch” decisions (a reward-by-transition interaction effect on subsequent first-stage choices). In our analysis, participants full trial-by-trial choice sequence in the task was fit with a computational reinforcement-learning model that gauges the degree to which participants choices are better described by a model-based or a model-free reinforcement learning algorithm.

**Figure 3:**
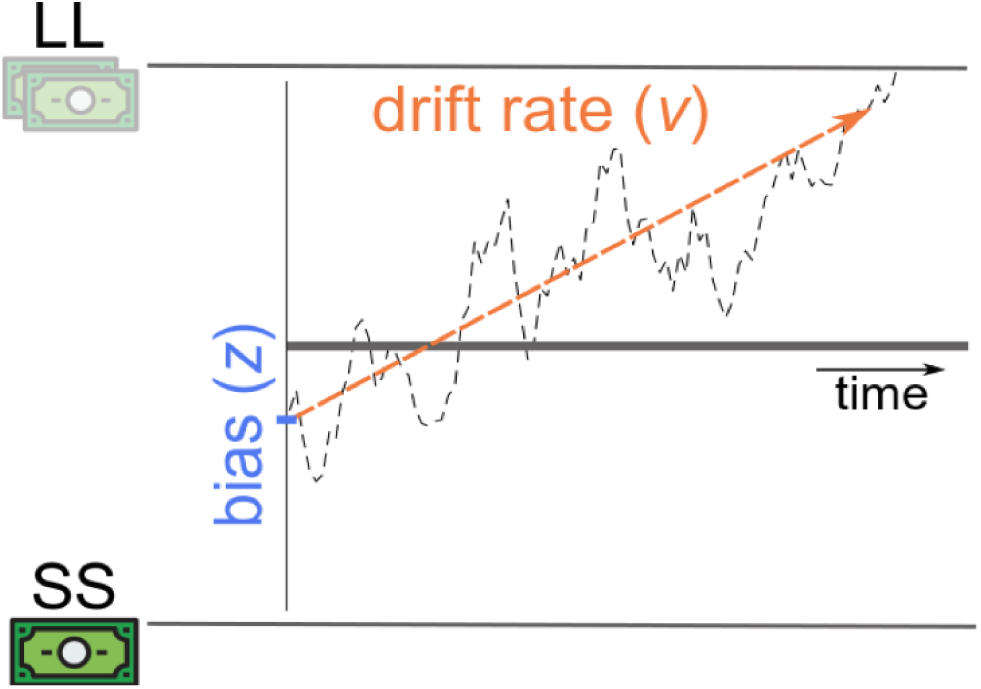
Decisions arise from a combination of bias z and drift rate v, which capture, respectively, the influence of reflexive and deliberative processing on choice. In this illustration, a choice for the LL (top) is disfavored by a static bias towards selecting SS (bottom). However, this bias is overcome during the course of deliberation, by evidence accumulated in favor of the LL at a rate (v) fast enough to reach the upper threshold before the deadline. Thus, the degree of “patience” displayed for a given choice can result from either a static, between-trials bias towards choosing LL, a deliberation of the values on the specific trial, or a combination of the two processes.

## 3 Computational Modeling

### 3.1 Intertemporal choice models

#### 3.1.1 Hyperbolic Model of Temporal Discounting

A discount rate was used as a summary of participant behavior. Discount rates (k) quantify the degree to which the subjective value of a future reward is discounted as a function of time, and can be estimated from a participants choices. While many functional characterizations of temporal discounting have been proposed, prior research suggests that intertemporal choice behavior is particularly well-described by a hyperbolic discount function [26] 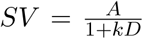, where the subjective value SV of a future monetary reward is calculated by dividing the objective amount A by one plus the discount rate k times delay D (here measured in days). Thus if the delay is 0, the subjective and objective amount are equal. With increasing delay time, or discount rate, the subjective value decreases. Parameters (discount rate, softmax temperature) were determined from the observed choices of each participant by using maximum likelihood iterative optimization (fminunc) in MATLAB (Mathworks, Natick, MA, USA).

#### 3.1.2 Drift-diffusion models (DDM)

To investigate whether future-oriented choices were made on the basis of deliberation, we implemented several variants of a canonical sequential sampling model, the drift-diffusion model (DDM; [11]). The models expressed different ways that the deliberative process might depend on the three elements of each choice: the value of the larger, later option (*V_LL_*), the value of the smaller, sooner option (*V_SS_*), and the delay to receipt of the LL (*T*). These values informed the decision process by setting the rate at which sampling proceeded toward the decision threshold (drift rate, or *v*) and/or the starting point of the accumulation process (bias, or *z*) 3.

Model parameters were specified as a hierarchical mixed-effects regression using the HDDM framework [27]. Each model set the v and/or z parameters as specified, with the remaining parameters (t, or the non-decision time, for all models, and sz, or the trial-by-trial variability in the starting point, for models that did not specify a trial-by-trial regressor for z) fit freely to each subject. Fitting was performed using a Markov-Chain Monte Carlo (MCMC) procedure, with chains set to 10000 total samples and 5000 burn-in samples.

The value difference model captured the hypothesis that choices were insensitive to T, by setting the drift rate as a function of *V_LL_* − *V_SS_*.

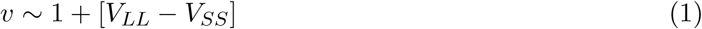

The bias-drift combination model captured the hypothesis that each subject used an idiosyncratic combination of instantaneous value difference and time-sensitive deliberation, by fitting separate coefficients for the starting point, as a function of value difference, and drift rate, as a function of (log) time to LL receipt.

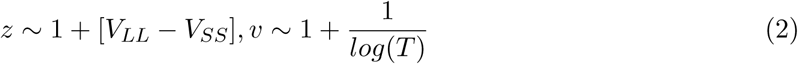

Our primary model of interest was the scaled difference model, which effectively jointly fit the discount parameter with the multiplicative coefficient component of the drift rate by setting drift equal to the reference-dependent value *V_LL_* − *V_SS_*, scaled by *log*(*T*):

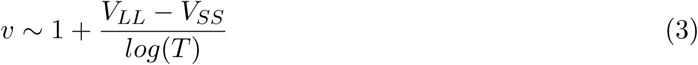

Comparing across types of models and tasks. These analyses gave us two families of models, with not only different numbers of parameters but also different types of input data choices for the hyperbolic model, choices and RT for the DDMs. To compare models on equal footing, we used leave-one-out cross-validation to assess the ability of the models to predict choices out-of-sample. For this analysis, we reduced each subjects dataset to their first 96 choices. Each model was then fit 96 times to populations in which all subjects performed the same sequence of 95 trials. This fit was evaluated by computing the choice probability (*p_s,m,i_*) that the model m would select as the subject s did on the held-out trial i. For DDM models, this choice probability was computed by generating 1,000 simulated choices using the parameters (*v, z, t, sz*) fit to the training set, and the settings (VLL, VSS, T) of the held-out trial, and computing ps,i as the fraction that resulted in the same choice as did the subject. For the hyperbolic model, *p_s,i_* was calculated exactly by using the fit discount parameter k and inverse softmax temperature. We computed the log-likelihood of each model m for each subject s by summing choice probabilities across test trials: *LL_m,s_* = −∑*_i_log*(*p_m,s,i_*).

Finally, the relative fits of these models were assessed using the Bayesian model selection procedure of [28], which takes model identity as a random effect. Bayes factors were computed on the basis of the per-subject log likelihood (choice probabilities), and models were compared using by submitting the log model evidences directly to the spm BMS.m routine from SPM8 (http://www.fil.ion.ucl.ac.uk/spm/software/spm8/).

All correlation statistics are Pearsons R, or in the case of partial correlations. When analyzing between-task correlations for the subset of subjects who performed the two-step task, robust correlations were performed after removal of bivariate outliers, using the MATLAB toolbox Corr toolbox [29]. In these cases, we report the 95% confidence interval (CI) of 1,000 bootstrap runs. All of the correlations reported here are robust to the exclusion of bivariate outliers, unless otherwise noted.

### 3.2 Two-step RL model

Indices of model-based learning in the two-step RL task were derived via Bayesian estimation using a variant of the computational model introduced in [23]. The model assumes choice behavior arises as a combination of model-free and model-based reinforcement learning. Each trial t begins with a first-stage choice *c*_1_*_,t_* followed by a transition to a second state *s_t_* where the participant makes a 2nd stage choice *c*_2_*_,t_* and receives reward *r_t_*. Upon receipt of reward *r_t_*, the expected value of the chosen 2nd stage action (the left vs. the right alien) 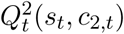 is updated in light of the reward received.

The model assumes 2nd stage choices arise due to a learned value function over states and choices *Q*^2^(*s, c*). The function updates the value for the chosen action by integrating the reward received on each trial using a simple delta rule,

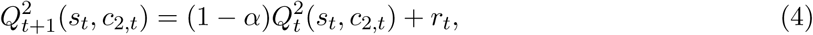

where *α* is a free learning rate parameter that dictates the extent to which value estimates are updated towards the received outcome on each trial. Readers from RL backgrounds will recognize that this update (Eq. 4) differs from the standard delta rule, *Q*(*s, a*) = (1−*α*)*Q*(*s, a*)+*αr*. In Eq. 4 and in similar references throughout, the learning rate *α* is omitted from the latter term. Effectively, this reformulation rescales the magnitudes of the rewards by a factor of 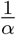 and the corresponding weighting (i.e., temperature) parameters *β* by *α*. The probability of choosing a particular 2nd stage action *c*_2_*_,t_* in state *s_t_* is approximated by a logistic softmax: 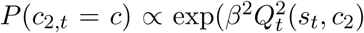 with free inverse temperature parameter *β*^2^ normalized over both options *c*_2_.

First-stage choices are modeled as determined by a combination of both model-free and model-based value predictions about the ultimate, 2nd stage value of each 1st stage choice. Model-based values are given by the learned values of the corresponding 2nd stage state, maximized over the two actions: 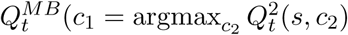, where *s* is the 2nd stage state predominantly produced by stage-1 choice *c*_1_. Model-free values are learned by two learning rules, TD(0) and TD(1), each of which updates according to a delta rule towards a different target. TD(0) backs-up the value of the stage-1 choice on the most recent trial 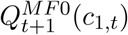 with the value of the state-action pair that immediately (i.e., lag-0) followed it: 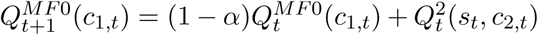, whereas TD(1) backs up its value estimate 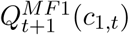 by looking an additional step ahead (i.e., lag-1) at the reward received at the end of the trial: 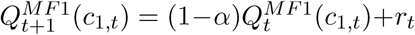. Stage-1 choice probabilities are then given by a logistic softmax, where the contribution of each value estimate is weighted by its own free temperature parameter: 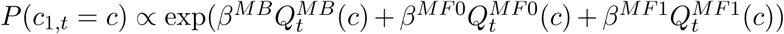. At the conclusion of each trial, the value estimates for all unchosen actions and unvisited states are decayed multiplicatively by *γ*, reflecting the assumption that value estimates decay exponentially by a rate of 1 − *γ* over successive trials. In total the model has six free parameters: four weights (*β*^2^*,β^M B^,β*^*M F*0^,*β*^*M F*1^), a learning rate *α*, and a decay rate *γ*.

This model makes two minor departures from some of the standard parameterizations outlined in prior work [23, 22]. First, whereas earlier models posit a single model-free choice weight *β_MF_* and use an eligibility trace parameter *γ* ∈ [0, 1] to control the relative contributions of TD(0) and TD(1) learning, here, as in recent work by [21], model-free valuation is split into its component TD(0) and TD(1) stages, each with separate sets of weights and Q values. The decay parameter *γ* represents an additional change in variables from other formulations. Although RL traditionally focuses on the process of updating the value of chosen actions (or state-action pairs) in light of their outcomes, substantial evidence from both theoretical and empirical work [30] suggests that the values of unchosen actions and unvisited states depreciate over time. Beyond this theoretical motivation, including a decay parameter accounts for the empirical observation that people tend to ‘stay (repeat) their most recent choice in a given stage; as the decay rate approaches 1, the value of the unchosen actions decrease towards zero, increasing the relative value of the chosen action on the next trial. By contrast, previous models [23] have operationalized choice perseveration using a “stickiness” parameter, which is implemented as a recency bonus or subjective “bump” in the value of whichever first stage action was chosen on the most recent trial (irrespective of reward).

The six free parameters of the model, *β*^2^, *β^M B^*, *β^M F^*^0^, *β^M F^*^1^, *α, γ*, were estimated by maximizing the likelihood of each individuals sequence of choices, jointly with group-level distributions over the entire population using an Expectation-Maximization procedure [31]. The resulting set of per-subject model-based weightings served as our primary index of model-based learning for further analysis of individual differences.

## 4 Results

### 4.1 Two paths to patience in intertemporal choice

Participants (N=494) made a series of binary decisions between two options one, a smaller reward, delivered sooner (SS), the other, a larger reward, to be delivered later (LL). On average, participants selected the patient option on 42.127% (0.645%) of trials. Response times to choose LL were longer, on average, than for choosing SS (across participants, mean RT(LL) = 1977.994 25.323ms; mean RT(SS) = 1831.7 ± 19.322ms; mean difference 146.294ms [95% CI: 111.528, 181.06] t(493)=8.268, p<.001).

The traditional approach to analyzing these intertemporal choice tasks is to measure the degree to which participants display a preference for early delivery of rewards by fitting a discount factor that decreases the subjective value of each option based on the time until receipt. We fit participant choices with a hyperbolic discounting model [6, 26]. The fit value of the hyperbolic discount factor k was 0.023 ± 0.002, with choice variability parameter of 0.291 ± 0.008.

We next fit an evidence accumulation model to participant choices and RT. Evidence accumulation algorithms are commonly implemented as models of binary value-based choice, with the decisiveness of deliberation the drift rate of evidence accumulation at each decision set to be a function of the difference between the subjective value of the two options. A higher value of the drift rate leads to faster choices and less-variable outcomes (Figure 3). Consistent with the idea that patience in the ITC task is linked to the rate of deliberation, more patient choosers were both faster to select the LL option (correlation between discount factor log(k) and mean RT to LL per participant: *R* = .304, *p* < .001) and, at the same time, less variable in their choices (correlation between discount factor log(k) and choice noise: *R* = −.361, *p* < .001).

To formally test whether such a mechanism could explain choices in our task, we fit several variants of a DDM to participant choices and reaction times; the variants differed in their specification of the relationship between drift rate and bias and the choice settings (*T_LL_,V_LL_,V_SS_*; see Methods for specifications). The best-fitting such model, the scaled-difference model, does not inherently discount option values, but instead produces stochastically-varying estimations with variance that increases with time to receipt.

We compared the best-fitting sampling model and the hyperbolic-time-preferences discounting model on their ability to predict held-out choices: For each of the N trials, the model parameters were fit to the other N-1 trials, and the fit model predicted the outcome for trial n. The likelihood is the sum, across all N fits, of the log choice probability of the held-out trial, thus allowing comparison of both models on equal footing. Consistent with the idea that these models capture the same behavior but with varying levels of specificity as to the mechanism, neither was a superior explanation of out-of-sample choices (average difference in log-likelihood: 2.526 ± 0.417; exceedance probability p=.633).

Two parameters of the sampling model capture qualitatively different, but non-exclusive, mechanisms by which an agent could choose LL: the bias z an offset that specifies the relative amount of evidence needed to select LL and the drift rate v the rate at which evidence is accumulated in favor of the higher-valued option (Figure 1). The first component can reflect simple rule-guided strategies such as “Choose LL when its value is greater than $50” or “Choose SS when the delay is greater than 10 days.” The second component reflects dynamic reasoning, for instance of the form “What will I do with $10 today?” or “What might I do with $100 in 50 days?” Regardless of implementation, the key distinction between the two lies at the computational level, where they capture different timescales of value arbitration: the bias reflects all processing whose influence is fixed before the presentation of the current options, whereas the drift rate captures deliberative processing that interacts with the specific conditions of the current choice (e.g. value of each option, and time until delivery).

Model parameters varied considerably between participants (Figure 4). As expected, the discount factor log(k) was reliably correlated with both drift rate v (R=-.852, p<.001) and bias z (R=-.517, p<.001). However, the v and z parameters were, themselves, correlated across participants (R=.401, p<.001). Supporting the hypothesis that separate processes influencing discounting, each parameters correlation with log(k) was reliable after controlling for the other (partial correlation of log(k) and v, controlling for z: *ρ*=-.823, p<.001; log(k) and z, controlling for v: *ρ*=-.365, p<.001) (Figure 5).

Given this decomposition, we examined in more detail the relationship between patience and choice noise. Consistent with the idea that the reduction in choice noise with patience is driven by the deliberative process and an increasing number of samples per unit time yielded more consistent value estimates partial correlation revealed a reliable relationship between choice noise and drift rate v when controlling for bias z (*ρ*=.361, p<.001), but not between and z when controlling for v (*ρ*=-.04, p=.375). This effect was reliable even after controlling for the overall relationship between choice noise and patience: was correlated with v when controlling for both log(k) and z (*ρ*=.133, p=.003). This analysis supports the hypothesis that choices result from dynamic option evaluation that involves successive samples of noisy evidence about option values.

In sum, we found that an evidence accumulation model allows us to decompose two paths to patience in intertemporal choice tasks. Specifically, while both drift rate v and bias z predict patient choices, captured by discount factor k, only v predicts deliberation of the sort that results in less-noisy choices. Next, we investigated whether this deliberative process was also related to a different type of choice behavior: planning in a sequential decision-making task.

**Figure 4:**
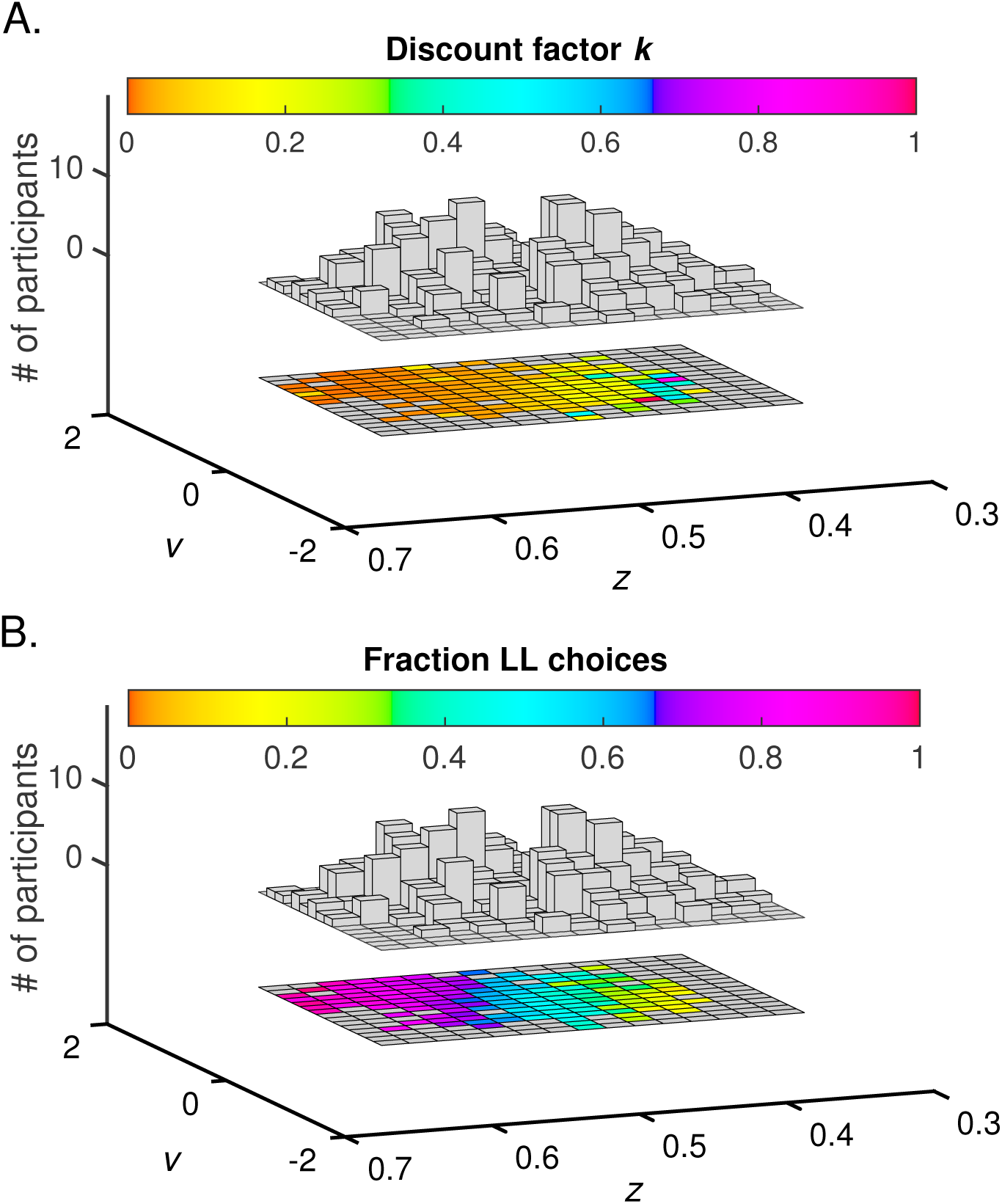
Drift rate and bias both contribute to patient choice. Choices in the intertemporal task, measured by the fraction of trials on which LL was chosen (A) and discount factor k (B), are a combined function of drift rate v and bias z (N=494). The discount factor log(k) was reliably correlated with both drift rate v (R=-.852, p<.001) and bias z (R=-.517, p<.001), but the v and z parameters were, themselves, correlated across participants (R=.401, p<.001). Two clusters of participants reflect different combinations of the two parameter values, yielding different proportions of patient choice.

**Figure 5:**
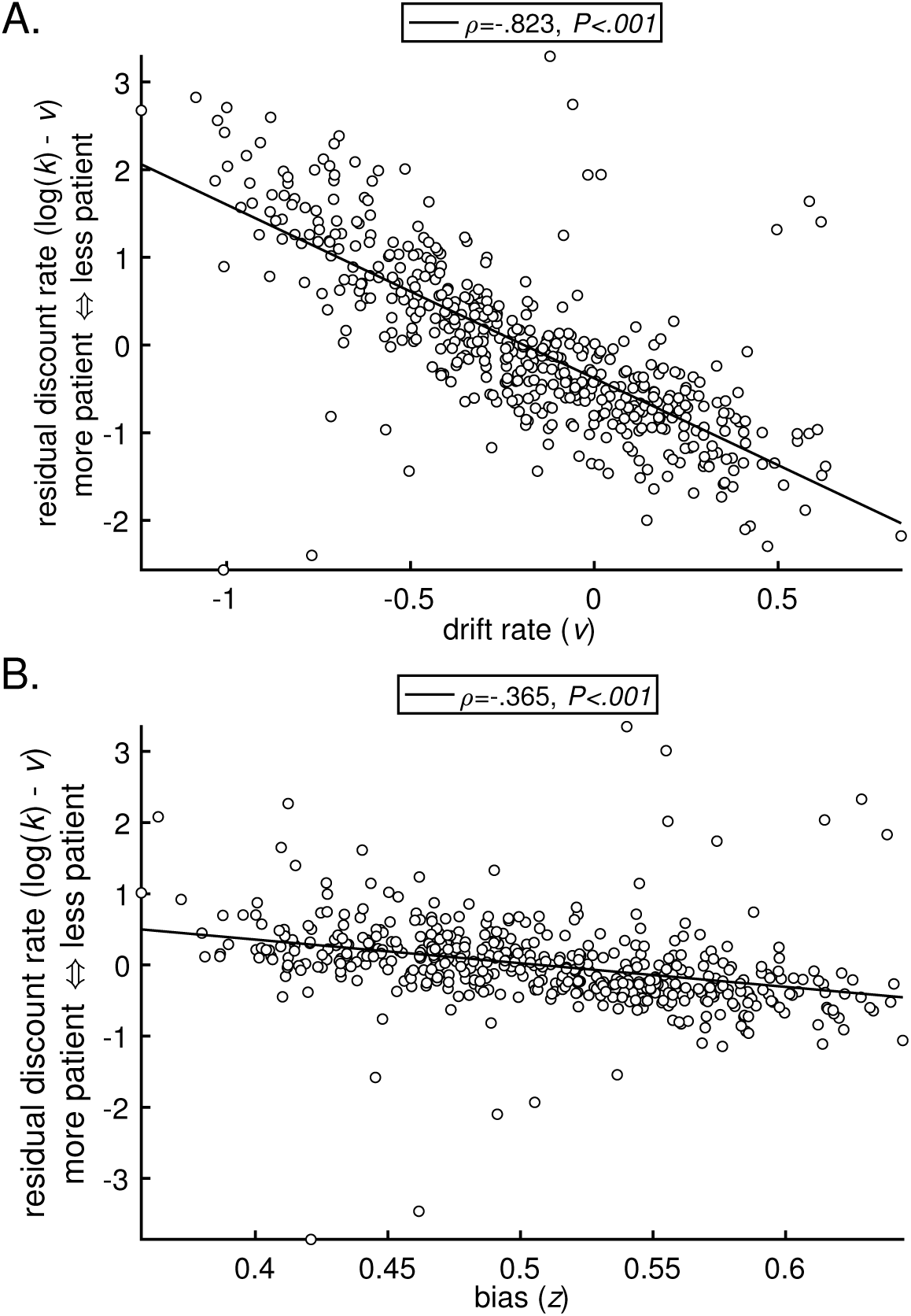
Hyperbolic discount factor is correlated with drift rate and bias. Hyperbolic discount factor (log(k)) is independently correlated with both the drift rate v (top) and bias z (bottom) of the DDM (N=494). Each parameters correlation with log(k) was reliable after controlling for the other (partial correlation of log(k) and v, controlling for z: *ρ*=-.823, p<.001; log(k) and z, controlling for v: *ρ*=-.365, p<.001).

### 4.2 Model-based planning is uniquely predicted by deliberative intertemporal choice

A subset of participants (N=32) performed a separate “two-step task” (Figure 1); [23, 24]). Following the standard approach to analyzing behavior in this task, we fit a computational reinforcement learning model to participants choices in the two-step task (see Methods). When participants properly learn (and use) the transition structure, and properly learn to credit rewards received after a “Rare” transition, their value of MB will be higher.

We first evaluated the relationship between behavior in the two tasks. Consistent with the idea that model-based planning indexes a general tendency towards forward-looking choice, participants with higher values of *β_MB_* were also more likely to choose the LL option (R(*β_MB_*, %LL)=.381, p=.015). Despite this, the model-based planning index *β_MB_* did not correlate with discount factor (R(*β_MB_*, log(k))=-.073, p=.346).

Although the relationship between model-based planning and patient choice was not mediated by the hyperbolic discount rate, we reasoned that such a relationship might be observable via the decomposition of discounting behavior that we identified previously, into drift rate v and bias z. Consistent with our hypothesis that a common deliberative mechanism underlies both intertemporal choice and model-based planning, only the correlation between MB and v was significant (*v*: R=.328, p=.033; *z*: R=.123, p=.251) (Figure 6). Confirming the specificity of this relationship, only v remained a significant predictor of MB when each correlation was taken while controlling for the contribution of the other parameter and log(k) (*v*: *ρ* =.374, p=.017; *z*: *ρ* =.007, p=.485) 6.

Taken together, these results demonstrate that apparent time preferences in intertemporal choice tasks arise from a process with two, computationally distinct components one a form of deliberation over the choice options, and the other a more automatic approach to binary choice. This decomposition of patient choice into two components reveals a relationship between intertemporal and model-based choice that is not observable in the standard model of discounted choice.

**Figure 6:**
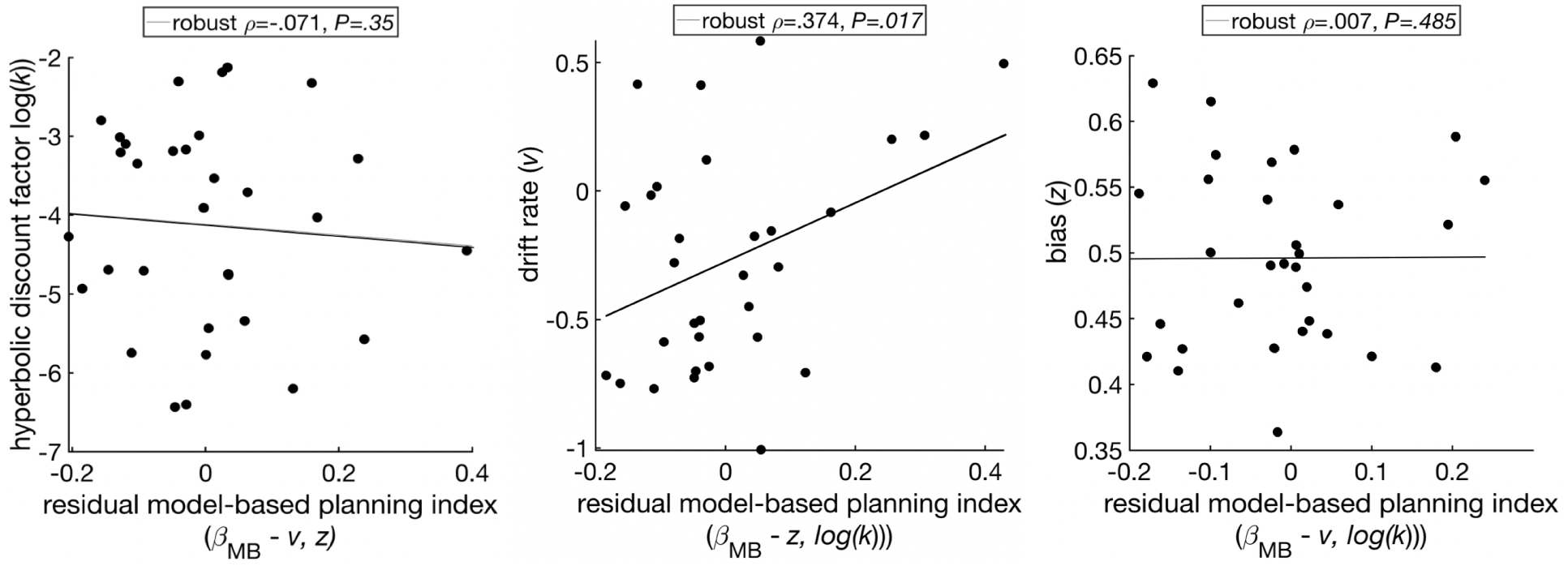
Model-based planning is correlated with drift rate, not discount factor or bias (N = 32). Despite not being correlated with hyperbolic discount factor (log(k), left), or DDM bias (*z*, right), model-based planning (MB) is correlated with DDM drift rate (*v*, middle) as fit to choices and RTs in the separately-administered intertemporal choice task. The correlation between MB and v is significant even after controlling for z and log(k) (robust =.374, [95% CI: 0.011, 0.671], middle).

## 5 Discussion

The ability to prospectively evaluate potential future outcomes is a critical feature of adaptive choice. Nearly every decision we face involves comparison between actions whose outcomes are not contemporaneously observable. In economics and psychology, this sort of decision has been studied as reflecting an idiosyncratic degree of preference for earlier consumption of rewards. However, a similar set of behaviors can arise without inherent time preference, but instead from a mechanism where evaluation of the choice options depends on the time until they are delivered. One such candidate mechanism is prospective evaluation of the sort proposed to underlie model-based planning in sequential reinforcement learning (RL) tasks. Here we found that individuals whose behavior in an RL task indicated increased recruitment of model-based learning strategy, which relies on prospective evaluation, also showed greater patience in a separate intertemporal choice (ITC) task. These behaviors corresponded at a computational level, and also an algorithmic one. We decomposed choices in the ITC task on the basis of their relative recruitment of reflexive or deliberative processes and found that the latter was exclusively associated with model-based planning.

Our finding may provide a unifying explanation of seemingly conflicting previous results about the external validity of the discounting measure, and in particular about the relationship between discount factors and measures of model-based planning [32, 33]. Specifically, the fact that patient choice can be decomposed into reflexive and deliberative components, each of which predicts the standard discount rate, raises the possibility that failures to find correspondence between discounting and other measures might be attributable to discounting that relies in larger part on the former, reflexive, process, rather than the more deliberative component. Such a reflexive approach to patient-seeming choice could arise from the fact that laboratory measures of intertemporal preferences generally involve repeated choices that, even if they are not sequential in nature, may support the adoption of cross-trial strategies tuned to the specific task at hand. Along these lines, a recent report that found no correlation between measured discounting and model-based planning [33] assessed discount factors using a staircasing procedure, which by its sequential nature may explicitly encourage across-trial bias-setting of the sort captured by the reflexive, but not deliberative, component. At the same time, the fact that these measures are positively correlated with each other suggests that there is not an explicit trade-off, and task features that encourage one need not diminish the other. Further work is necessary to understand what types of tasks encourage different evaluation strategies, and to what degree each corresponds to other behaviors and personality traits traditionally associated with patience in intertemporal choice.

These results concord with recent work showing that temporal discounting does not require a preference over temporal delivery. [10], as long as the forecasts generated by the mechanism are more variable when predicting outcomes farther away in time. They further showed that hyperbolic discounting describes the behavior of this rational agent when forecast variance is linearly increasing with time. In other words, the degree of (im)patience corresponds to the tradeoff between the potential of a larger reward, and the internal cost of reliably simulating that reward. Such a tradeoff parallels analyses showing that the balance between model-based and model-free decisions result from a cost/benefit or speed/accuracy tradeoff [18].

In the brain, the relative influence of model-based vs. model-free valuation is negotiated via the dynamic [18]. The planning computations that support model-based behavior have been shown to involve contributions from dorsolateral prefrontal cortex, dorsomedial striatum, and hippocampus [19, 20]. Patient intertemporal choice is also thought to involve a search through future outcomes, and has been shown to engage a similar cortico-subcortical network of brain regions including the hippocampus [34, 5] and dorsolateral prefrontal cortex [34]. Further support for the role of prospective simulation in patient choice is given by evidence linking numerous cognitive processes related to episodic memory, such as self-projection, episodic future thinking, mental time travel, and scene construction, to simulating the outcomes of hypothetical actions [35, 36, 37, 38].

Broadly, the computational flexibility of the RL framework offers future research a viable tool for dissociating sub-mechanisms of intertemporal choice. While such mechanistic distinctions are always important when trying to relate behavioral differences to neural substrates, the importance of untangling the many potential causes of impatient choice is underscored by the fact that observations of impulsivity across multiple clinical populations are likely to arise from different causes. Recovery from substance abuse, for example, is thought to rely on the ability to take the perspective of your future self in order to envision the long-term benefits of sobriety in spite of its tension with the contemporaneous drug craving. While many treatments focus on building proper habits [39], recent findings support the promotion of prospective future simulation as an effective approach for decreasing drug consumption [40]. A computational framework that reliably distinguishes multiple paths to patience could provide theoretical grounding for when to apply each approach, and help therapists take into account the myriad environmental and personal factors known to alter deliberative decision-making under uncertainty.

## Acknowledgements

This work was supported by NIH Grant R03DA038701, NSF CAREER grant 1654393 and a generous gift from the Mortimer D. Sackler MD family.

